# DeLTA 2.0: A deep learning pipeline for quantifying single-cell spatial and temporal dynamics

**DOI:** 10.1101/2021.08.10.455795

**Authors:** Owen M. O’Connor, Razan N. Alnahhas, Jean-Baptiste Lugagne, Mary J. Dunlop

## Abstract

Improvements in microscopy software and hardware have dramatically increased the pace of image acquisition, making analysis a major bottleneck in generating quantitative, single-cell data. Although tools for segmenting and tracking bacteria within time-lapse images exist, most require human input, are specialized to the experimental set up, or lack accuracy. Here, we introduce DeLTA 2.0, a purely Python workflow that can rapidly and accurately analyze single cells on two-dimensional surfaces to quantify gene expression and cell growth. The algorithm uses deep convolutional neural networks to extract single-cell information from time-lapse images, requiring no human input after training. DeLTA 2.0 retains all the functionality of the original version, which was optimized for bacteria growing in the mother machine microfluidic device, but extends results to two-dimensional growth environments. Two-dimensional environments represent an important class of data because they are more straightforward to implement experimentally, they offer the potential for studies using co-cultures of cells, and they can be used to quantify spatial effects and multi-generational phenomena. However, segmentation and tracking are significantly more challenging tasks in two-dimensions due to exponential increases in the number of cells that must be tracked. To showcase this new functionality, we analyze mixed populations of antibiotic resistant and susceptible cells, and also track pole age and growth rate across generations. In addition to the two-dimensional capabilities, we also introduce several major improvements to the code that increase accessibility, including the ability to accept many standard microscopy file formats and arbitrary image sizes as inputs. DeLTA 2.0 is rapid, with run times of less than 10 minutes for complete movies with hundreds of cells, and is highly accurate, with error rates around 1%, making it a powerful tool for analyzing time-lapse microscopy data.

**Author Summary:** Time-lapse microscopy can generate large image datasets which track single-cell properties like gene expression or growth rate over time. Deep learning tools are very useful for analyzing these data and can identify the location of cells and track their position over time. In this work, we introduce a new version of our Deep Learning for Time-lapse Analysis (DeLTA) software, which includes the ability to robustly segment and track bacteria that are growing in two dimensions, such as on agarose pads or within microfluidic environments. This capability is essential for experiments where spatial and positional effects are important, such as conditions with microbial co-cultures, cell-to-cell interactions, or spatial patterning. The software also tracks pole age and can be used to analyze replicative aging. These new features join other improvements, such as the ability to work directly with many common microscope file formats. DeLTA 2.0 can reliably track hundreds of cells with low error rates, making it an ideal tool for high throughput analysis of microscopy data.

## Introduction

The automation of hardware and software for microscopy has resulted in researchers’ ability to generate massive datasets containing images of cells over time. For example, in a recent high throughput experiment Bakshi *et al*. imaged 10^8^ *Escherichia coli* over days by acquiring 705 field of views every few minutes (1). Additionally, recent studies have used closed-loop microscopy and optogenetic platforms to control gene expression in single cells in real time (2–4). These improvements in microscopy have motivated the need for automated image analysis, as traditional approaches that require manual error correction cannot keep pace with the size of these new datasets or the rate at which they can be acquired.

To address this, researchers need tools that are rapid, accurate, and require minimal input from the user. This combination of needs is well suited for deep learning-based approaches, and deep convolutional neural networks have enabled fast and accurate analysis of images. Specifically, the U-Net architecture has emerged as the state-of-the-art convolutional neural network for biomedical applications (5). U-Net uses a “U”-shaped network architecture with a contraction path, where successive convolutional layers are progressively down-sampled, followed by a symmetric expansion path where the low-resolution but high-level encoding of the input image is up-sampled back to the original resolution. This approach has been widely successful for segmentation of cells (6–10) and for tracking cells from frame-to-frame within time-lapse images (7,9).

Here, we focus on analysis of bacterial time-lapse microscopy data growing in two-dimensional settings such as agarose pads. With rapid cell cycle times, small cell sizes, and high throughput microfluidic devices, it is possible for researchers to generate large datasets containing thousands of single cells over periods of hours or days. As a result, researchers can use statistical analysis to study the subtle and complex effects of cell-to-cell heterogeneity, gene expression dynamics, and cell-to-cell interactions in isogenic populations (1,11). This has led to fundamental discoveries related to antibiotic resistance (12,13), and has allowed for accurate characterization of genetic parts and circuits (14). Further, single-cell time-lapse analysis has revealed that cell division in *E. coli* is asymmetrical, where daughter cells receiving the ‘old’ pole grow more slowly than daughter cells receiving the ‘new’ pole (15). However, these effects are subtle, necessitating measurements of many division events to determine statistically significant effects (16). Until recently, such studies necessitated painstaking semi-manual analysis and curation of microscopy data. Software based on traditional image analysis techniques such as Schnitzcells (17), Oufti (18), or SuperSegger (19) all require significant user input and post-processing. A few recent studies have proposed deep learning models for bacterial or yeast cell segmentation (6,8,10,20), however to our knowledge there is no integrated segmentation and tracking pipeline for two-dimensional timelapse analysis of bacteria.

In previous work, we developed the Deep Learning for Time-lapse Analysis (DeLTA) pipeline to analyze single-cell growth and gene expression in microscopy images (7). DeLTA uses two instances of the U-Net model to segment and then track cells. This allows for rapid and robust analysis of time-lapse movies. The original version of DeLTA focused on segmentation and tracking of cells in the ‘mother machine’ microfluidic device (21) where bacteria are constrained to narrow chambers where they grow in a single file line. This powerful design simplifies image analysis and enables experiments that run for many hours or days. However, this constrained geometry is not well suited for the study of two-dimensional effects such as diffusion of chemical signals or proximity-based effects and co-cultures with mixed populations of cells. Two-dimensional configurations, ranging from microcolonies growing on agarose pads to microfluidic growth chambers, can be used to measure spatial dynamics of cell-to-cell interactions. Examples include quorum sensing (22) and the effect of efflux pumps on neighboring cells in the presence of antibiotics (23). Segmenting and tracking cells in two dimensions are more challenging than for cells constrained within the mother machine. Segmentation becomes more difficult as microcolonies grow because images can contain hundreds of cells, where any given cell may have neighbors on all sides. The complexity associated with tracking also increases dramatically. In contrast to mother machine data, where frame-to-frame assignments can be limited to the small number of cells within the chamber (typically <10 cells), two-dimensional environments need to consider hundreds of possible assignments. Further, cells can move in any direction and may move large distances, for example if there is drift over the course of the movie.

In this manuscript we introduce DeLTA 2.0, a new version of DeLTA that segments and tracks cells in two dimensions. DeLTA 2.0 retains all the functionality of the original version and is fully compatible with mother machine data. The new version is available open source and uses a fully Python implementation. We have also introduced other improvements to the code to increase accessibility, such as the ability to work with images of arbitrary size and to accept many common microscope file formats as inputs. We show that DeLTA 2.0 can segment and track co-cultures of *E. coli* growing on an agarose pad. In addition to fluorescence and cell length, DeLTA 2.0 also records pole age. We use this to record replicative aging and compare the growth rate across generations. DeLTA 2.0 is a powerful tool for rapidly and accurately segmenting and tracking single cells on two-dimensional surfaces. It performs well on crowded images and requires no human intervention.

## Results

DeLTA 2.0 can track cell length, growth rate, fluorescence, and progeny over time for cells growing in a two-dimensional microcolony. The algorithm takes microscopy images as inputs. It then uses two U-Net convolutional neural network models, one for segmentation and one for tracking, and outputs single-cell data (Fig 1A). Segmentation generates information about cell morphology and can be used to identify both normal and filamented cells (Fig 1B-C). The tracking step reliably tracks cells across frames, recording division events when they occur (Fig 1D). The algorithm outputs single-cell resolution information for all cells within a field of view (Fig 1E). No human intervention is required to specify any input parameters. This is in sharp contrast to other methods, which typically require inputs, such as cell size or cell type (6,17–19,24).

**Fig 1.**
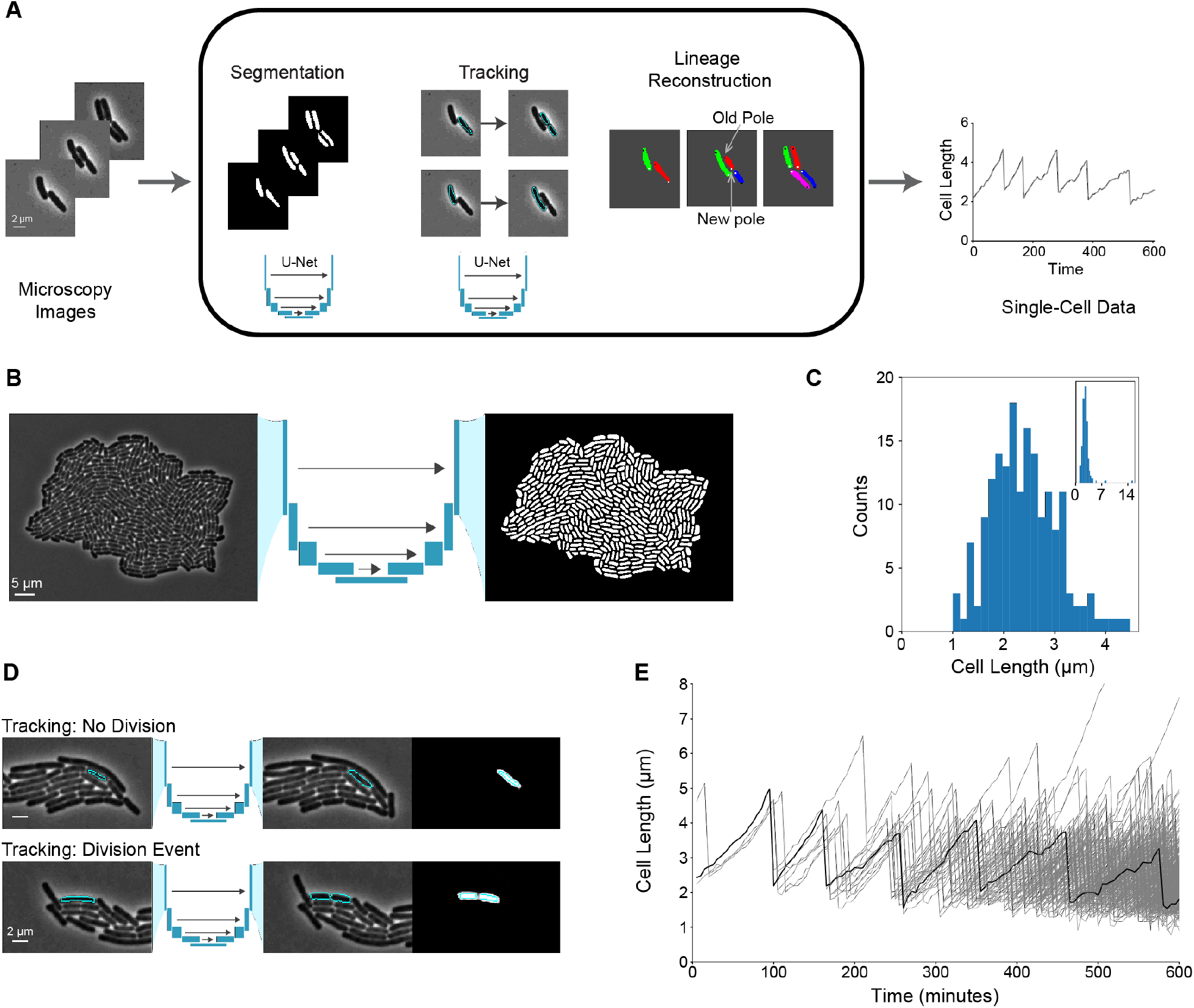
Segmentation and tracking of cells within a microcolony. **(A)** Workflow of DeLTA showing segmentation, tracking, and lineage reconstruction. **(B)** Phase contrast image containing an *E. coli* microcolony, which is input into a U-Net convolutional neural network to obtain segmentation results. **(C)** Histogram of cell lengths. Inset shows a zoomed out version with outliers included. **(D)** Cell tracking between frames. Representative examples of cell tracking with and without division are shown with a phase contrast image of the ‘previous frame’ on the left, a phase contrast image of the ‘current frame’ in the middle, and a grayscale image of the ‘prediction’ on the right. The ‘current frame’ shows the tracking prediction overlayed. The ‘prediction’ shows the U-Net output with the ground truth overlayed. **(E)** Plot of cell lengths over time. Black line is a representative example of one cell’s length as it grows and divides; all cells in the microcolony are shown in gray.

DeLTA 2.0 can process datasets of any dimension quickly and robustly. To evaluate its speed and accuracy, we used a movie from the literature that has been segmented and tracked with manual correction that DeLTA had not been trained on (Methods). It took 8 mins and 49 secs to conduct complete analysis of this time-lapse movie containing 69 frames, where the final frame contains 232 *E. coli* cells (S1 Movie). This analysis was conducted on a desktop computer with a NVidia Quadro P4000 graphics card. In addition to being fast, DeLTA also has a low error rate. Out of the 3,286 cells segmented in the test set, there was an error rate of 0.01%. We defined a correct segmentation prediction as any case where the cell annotation in the model prediction had more than three quarters of its pixels overlapping with the ground truth data. This allows subtle differences between the prediction and ground truth to be considered acceptable, such as slight discrepancies in the exact location of the cell perimeter. In addition, if the model made a prediction where it erroneously connected two cells together, we defined this as two errors. Erroneously predicting a split cell was counted as one error. Because it can be difficult to assess the exact frame at which a cell divides, we did not count predictions where the model determined division events to be up to three frames later or earlier than in the ground truth as an error. To assess the tracking error rate, we processed 6 movies of *E. coli* growing on agarose pads that the model had not been trained on. Out of the 17,622 tracking events, we measured an average error rate of 1.02%. We defined a correct prediction as a case where the cell assignment from one timepoint to the next matched the ground truth. Cases where the model assigned two cells as the daughters of a cell from the previous frame when there was in fact no division event, and cases where the model assigned no cell when the cell was in fact still within the field of view, were counted as tracking errors.

The DeLTA 2.0 algorithm has several improvements over the original version of DeLTA (7). The new code is a purely Python workflow; movies do not need to be pre- and post-processed in Matlab. This transition allows the entire pipeline to exist in an open-source framework. We do provide code that can be used to convert the output to a Matlab file for users that are more comfortable working in this environment for post-processing data. In addition, in DeLTA 2.0 we take advantage of the Bio-Formats toolbox for Python (25,26). This allows users to work directly with images in many common formats that are output via microscopy software, including nd2, czi, ome-tiff files, and many more without the need for any preformatting. We also made updates to the code that increased its flexibility, while optimizing for performance. For example, DeLTA 2.0 can accept input images of any size. For large images (>512×512 pixels), DeLTA 2.0 will automatically crop the image into smaller windows for segmentation and then stitch the outputs back together.

Since the original DeLTA code was optimized for images from the mother machine, where cells are constrained to one-dimensional chambers, tracking was relatively straightforward. In two dimensions, tracking is a more complex task and the number of cells that need to be tracked simultaneously increases dramatically. It can be challenging to identify which cells are associated with which lineage. To improve tracking speed, we crop a 256×256 pixel area around the cell of interest. This approach works because a single cell is expected to remain in the immediate vicinity of where it was in the preceding frame, so it is reasonable to restrict the search for daughter cells to the local area. These coordinates are then used to crop the three other inputs (previous phase contrast image, current phase contrast image, and current segmentation). The dimensions of these cropping windows can be changed in the configuration file.

To reduce overfitting, DeLTA 2.0 uses several new data augmentation operations while training. In addition to operations such as random shifting, scaling, rotation, flipping, and illumination which were present in the original software, we added three new functions, two for segmentation and one for tracking. To help deal with an occasional out of focus frame, we added a blurring function that slides a Gaussian kernel over the image (Methods). In addition, electronic noise is another issue when dealing with biological samples where the minimization of the total exposure to excitation light decreases the signal-to-noise ratio of the camera’s sensor. To deal with this, we added a function that adds Gaussian noise (Methods). To simulate exaggerated cell movement during tracking, such as when an agarose pad dries out and causes the field of view to shift over time, we wrote a new augmentation function that introduces image translations between different timepoints (Methods). These operations help expand the training dataset and allow the model to generalize to realistic conditions.

To showcase the utility of DeLTA 2.0, we performed several experiments where we grew *E. coli* microcolonies on agarose pads and analyzed the output. First, we used DeLTA 2.0 to distinguish differences in growth rate between antibiotic resistant and susceptible cells grown in the same field of view. In this experiment, we mixed two strains of *E. coli*, one containing a tetracycline resistance gene and a constitutively expressed red fluorescent protein (RFP) reporter, and the other without the resistance gene and containing a green fluorescent protein (GFP) reporter. We grew cells in a co-culture on an agarose pad containing an inhibitory concentration of tetracycline (0.5 μg/ml). DeLTA 2.0 reliably segmented cells within the image (Fig 2A). The tetracycline resistance gene allowed the RFP-expressing cells to grow well whereas the tetracycline sensitive GFP-expressing cells grew very slowly (Fig 2B). The RFP and GFP fluorescence of individual cells can be plotted over time and show the two distinct strains (Fig 2C). By extracting mean fluorescence levels for all cells within the time-lapse images, we found that fluorescence levels for the two populations were well-separated and maintained over time, as would be expected for the constitutive reporters. We also used DeLTA 2.0 to calculate the individual cell growth rates with respect to fluorescence. We observed two distinct clusters, corresponding to RFP cells that grew normally and GFP cells that grew slowly or did not grow (Fig 2D). These results highlight the ability to track cells with different properties simultaneously within the same image.

**Fig 2.**
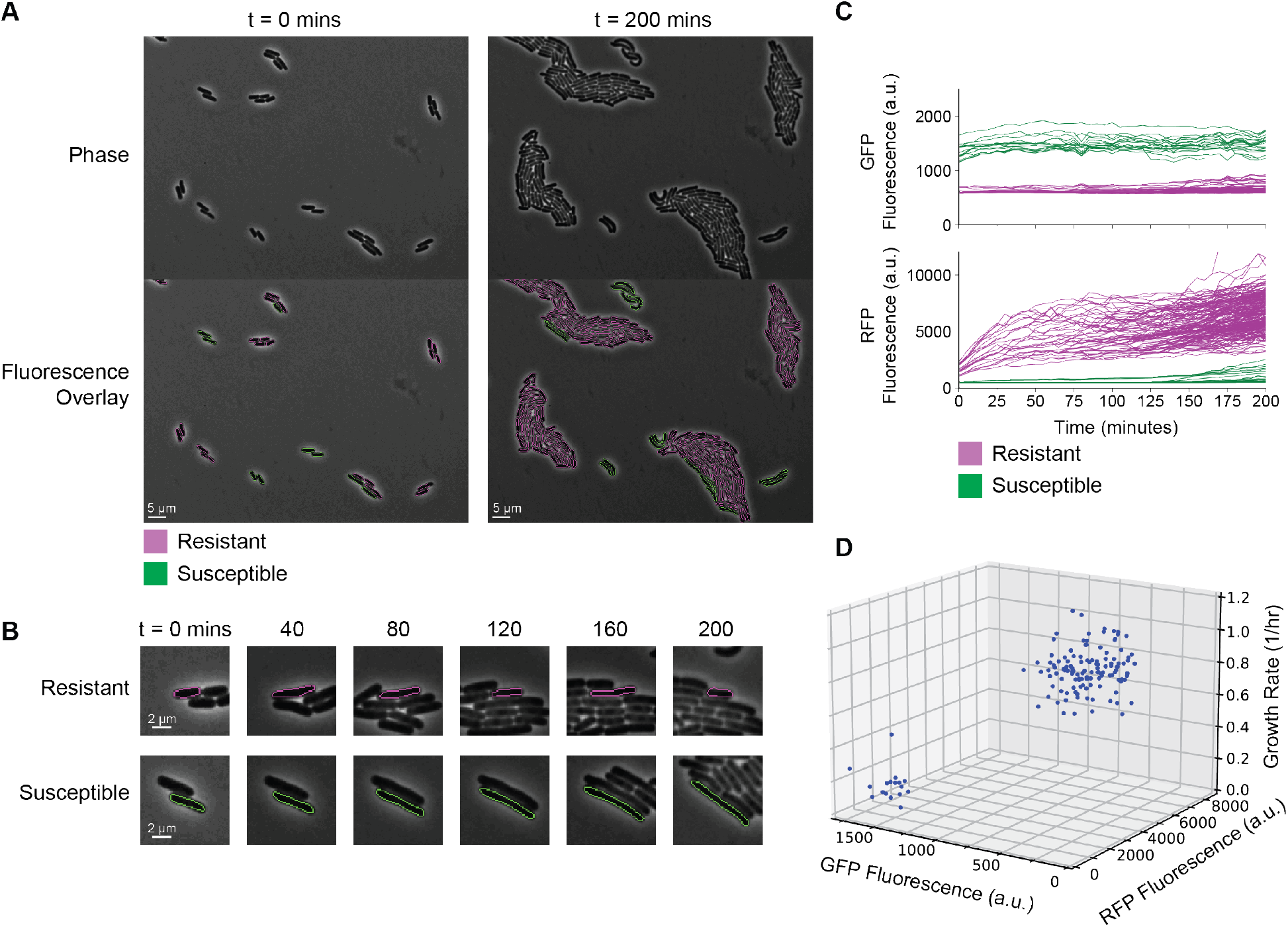
Resistant and susceptible strains of *E. coli* on agarose pads containing an inhibitory concentration of tetracycline. **(A)** Phase contrast images and associated fluorescence overlays. RFP expressing cells contain a tetracycline resistance gene and GFP expressing cells do not. The magenta and green outlines in the fluorescence overlay represent the resistant and susceptible cells, respectively. **(B)** Representative examples of antibiotic resistant and susceptible cells tracked over time. **(C)** RFP and GFP fluorescence tracked for individual cells over time. **(D)** GFP fluorescence versus RFP fluorescence for single cells plotted against growth rate. Fluorescence values are the averages over all the frames for that cell. For growth rate calculations, only cells that were present at t = 150 min were tracked, which is a time point mid-to-late in the movie. The analysis omits those cells that enter the field of view after t = 150 min since the growth rates become noisier with less data. Three outliers are omitted from this view to highlight differences in growth rates; all data points are shown in S1 Fig.

DeLTA 2.0 is well suited for measuring growth and gene expression for many cells within an image. Because of this, another potential application is the study of replicative aging within bacterial microcolonies. Recently studies have shown that non-genetic differences may be passed down to offspring, causing a modest but measurable change in growth rate (15,16,28–30). This can be tracked by recording pole age over time. Rod shaped bacteria have two poles, where one end of the cell is referred to as the ‘old’ pole if it was passed down from the mother. The pole formed after division is referred to as the ‘new’ pole (Fig 3A).

**Fig 3.**
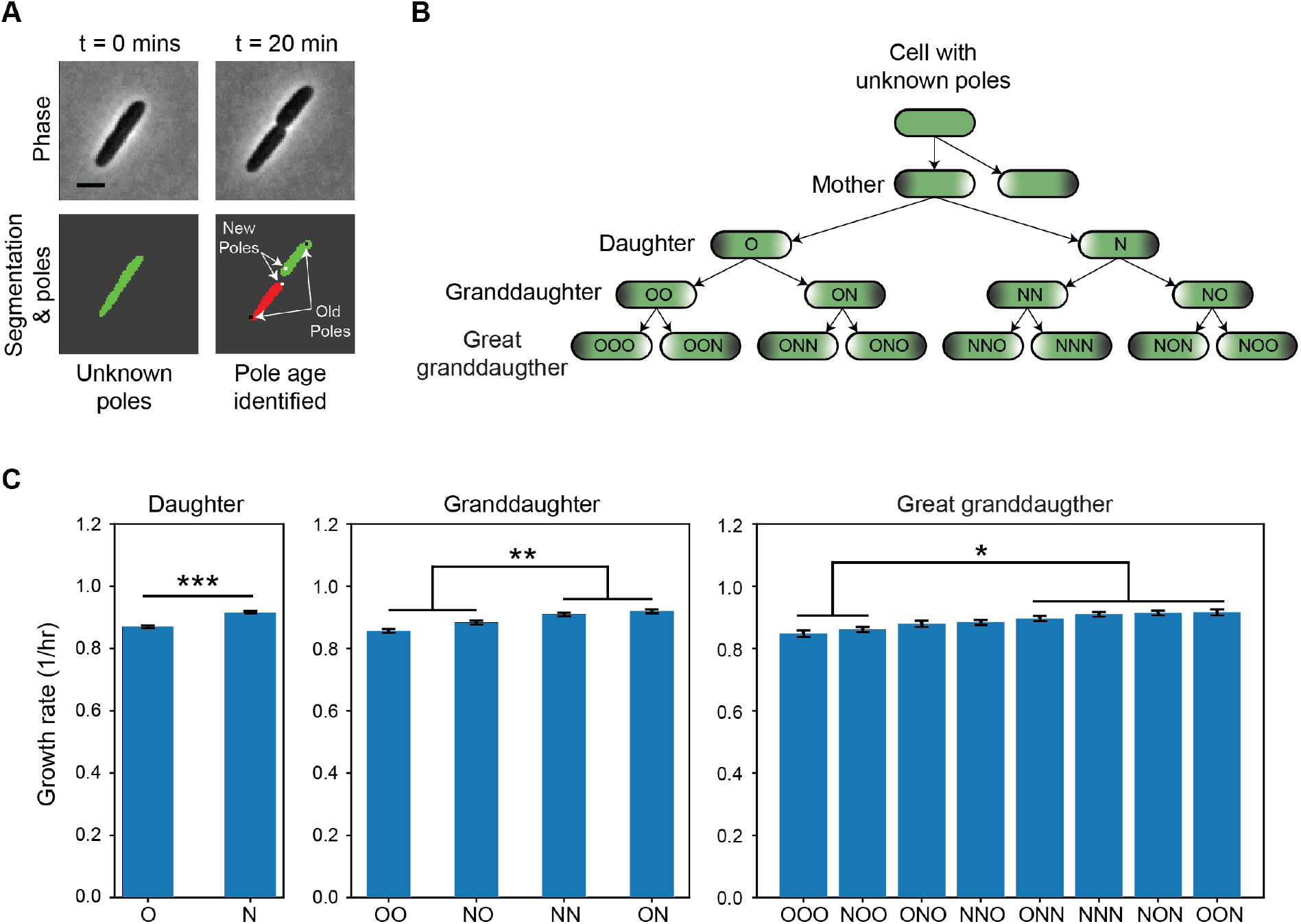
Pole age and its impact on growth rate. **(A)** Schematic showing how poles are passed down during a division. When a cell divides, the newly formed poles are defined as the ‘new’ poles (white dot) whereas the poles that were passed down from the mother are defined as the ‘old’ poles (black dot). Scale bar, 2 μm. **(B)** Pole assignment schematic. When the mother cell with known poles divides, the daughter cell that inherits the mother’s old pole is denoted ‘O’ whereas the daughter that inherits the mother’s new pole is ‘N.’ For each generation, either an O or an N is appended to the pole history. **(C)** Growth rate within each generation. The growth rate of an individual cell is calculated for the period right after the mother’s division until right before the cell divides again. To reduce noise, only cells present for at least three frames were included in the analysis. Daughters (n = 11,246 cells; two tailed unpaired t-test; p-value *** ≤ 0.001), granddaughters (n = 10,726 cells; one-way ANOVA test performed; p-value ** ≤ 0.01), and great granddaughters (n = 10,217 cells; one-way ANOVA performed; p-value *≤ 0.05). Error bars show standard error of the mean.

To date, many experiments studying pole age have been conducted in the mother machine microfluidic device due to the ease of tracking cells (15,21,30). However, a limitation of this approach is that it is only possible to track cells for a small number of generations because older generations are swept out of the imaging chamber while the mother cell’s old pole stays at the dead end of the chamber. For this reason, the original DeLTA algorithm did not track pole age information, and in a division event the algorithm simply assigned the cell closest to the dead end of the chamber to be the mother and the other cell to be the daughter. However, on two-dimensional surfaces cells can be aligned in any orientation, therefore DeLTA 2.0 assigns old and new poles after division based on the position of the septum. To highlight the ability to track pole age over time, we analyzed a movie of *E. coli* growing in unstressed conditions. At t = 0, we do not know the history of the cells, so the poles are initially unassigned (Fig 3B). After division, the mother’s poles that are passed down become the ‘old’ poles and the newly divided poles are the ‘new’ poles of the daughters. This proceeds over subsequent generations. To keep track of this, we denote the daughter that receives the old pole as ‘O’ and the new pole as ‘N’. When these daughter cells divide, they form cells that are granddaughters of the original mother cell. We append an O or N at the end of the pole sequence to record this. For example, the cell that inherits the original old pole is denoted OO and its sibling is ON. This continues through the generations, such that great granddaughters of the original mother have three letters in their pole sequence (e.g. OON).

First, we compared the growth rate of the old and new pole daughters from time-lapse movies of *E. coli* growing in unstressed conditions. Consistent with prior literature (15,16,28), we found that old pole (O) daughters grew more slowly than the new pole (N) daughters (Fig 3C). Next, we used the rich generational information provided by DeLTA 2.0 to test for differences in growth rate between granddaughters and great granddaughters with different pole ages. We found that growth rate differences were dependent on which pole the cell most recently received. For instance, OO and NO had growth rates that were lower than the NN and ON (Fig 3C). Therefore, the old versus new pole influence upon growth rate is dominated by effects that extend back only one generation. This result was consistent with great granddaughters as well, where growth rates tended to be slower in cells that most recently received an old pole (OOO, NOO, ONO, NNO) than in those that received a new pole (ONN, NNN, NON, OON). These results demonstrate how pole age information can be tracked with DeLTA 2.0, enabling studies on replicative aging.

## Discussion

In this work, we developed a deep learning pipeline that can process time-lapse images of bacterial microcolonies and output single-cell data. Utilizing the U-Net convolutional neural network architecture, the model can rapidly segment and track cells frame-to-frame with a low error rate. We applied this to successfully differentiate growth rates between sensitive and resistant strains of *E. coli* growing on an agarose pad. This demonstrates DeLTA’s new ability to measure co-culture dynamics, which are hard to capture in devices like the mother machine. DeLTA 2.0 retains all the functionality of the original version of DeLTA and can now be used on mother machine data or microcolonies of bacteria growing in two dimensions.

The current model works very well on standard rod-shaped bacteria such as *E. coli*, suggesting that analysis of cells with similar morphologies such as *Bacillus, Pseudomonas*, and *Salmonella* species will be straightforward. However, DeLTA 2.0 may require further training for cases where the appearance of the cells deviates from the data it was trained on. For example, although the model can handle some elongated cells (Fig 1B-C), it is not currently optimized for cells with highly filamented morphologies or cells undergoing stress. In addition, if the position of the cells within the field of view shifts dramatically from frame to frame, the current software does not perform well. This problem is exacerbated in conditions where the frame is crowded with cells. Thus, optimizing experimental conditions to minimize drift is important for high quality analysis.

Although segmentation works efficiently and has a low error rate, the model sometimes makes incorrect predictions about distinct cells being connected (S2 Fig). Within isolated images, these errors can be difficult to catch, even for humans. However, by looking at earlier and later frames it is usually possible to identify such errors because cells cannot divide and then merge back together. In the future, deep learning architectures that use time-series information such as recurrent neural networks could be combined with our models to improve segmentation by incorporating temporal context.

The tracking model runs robustly but can slow down when there are hundreds of cells in the image. Because every cell in the frame creates an input for the tracking model, this increases exponentially as the bacterial microcolony grows. Future efforts to optimize the tracking algorithm could help to address this by avoiding methods that scale linearly with the number of cells. In addition, initial tests suggest that it may be possible to decrease the size of the convolutional neural network by removing layers, though this would likely need to be customized for specific applications (S3 Fig). This could be applied to the segmentation or tracking model.

Overall, DeLTA can now process two-dimensional movies accurately and capture spatial dynamics in a high throughput manner with no human intervention. It works with many common microscopy file formats and extracts single-cell features such as cell poles, length, lineage, and fluorescence levels automatically and saves data into Python and Matlab compatible formats. As many microbiology researchers work with these types of data, we envision that this software can be used to increase the throughput of microscopy image analysis.

## Methods

### Implementation and network architecture

The U-Nets are implemented in TensorFlow/Keras. The code, installation instructions, and datasets are available open source on Gitlab: https://gitlab.com/dunloplab/delta. We developed a fully-integrated pipeline that can compile single-cell data from Bio-Formats compatible files or TIFF image sequences, but we also provide simpler scripts and data that illustrate the core principles of our algorithm for easy adaptation to different use cases. Network architecture for the segmentation and tracking model is as described in the original DeLTA publication (7), with modifications described as follows. In original DeLTA (7), the tracking model uses a softmax function as the final activation layer and a categorical cross-entropy loss function to produce three grayscale output images with 1’s in each layer representing the mother cell, daughter cell, and background, respectively. In DeLTA 2.0, the tracking model uses a sigmoid function as the final activation layer and a pixelwise-weighted binary cross-entropy loss function to produce a single grayscale output image with 1’s representing tracked cells (mother and potential daughter) and 0’s representing the background and the cells that did not track to the input cell.

### Loss functions and training

To train both models, we implemented a pixelwise-weighted binary cross-entropy loss function, as in the original U-Net paper (5). The loss function determines the difference between the model output and the ground truth which is then used to update the weights within the model. As in Ronneberger *et al*. (5), our loss function takes a weight map as an extra input to assign more importance to certain pixels in the ground truth during training. We used custom weight maps to improve segmentation on rod-shaped bacteria by increasing weights for the center of the cells and the borders between the cells. We also minimize weight on the background, where background is defined as anything in the image that is not a cell or border (S4 Fig). More specifically, we maximized the weights for the skeletons of the cells and borders, which are pixel-wide representations of binary objects in images (31). Determining the exact borders of the cells by eye is hard and partially arbitrary. To prevent the model from learning these arbitrary cell-border interfaces, we reduced the weights in these areas (S4 Fig). In addition, the background weights were set to be variable, where the weight increased with respect to an incorrect prediction (S5 Fig). The model outputs a number between 0 and 1, with 0 representing the background and 1 representing a cell. Since we had high confidence in the ground truths for the background, we were able to set the values of the weight map for the background to be equal to the actual prediction for the background. If the model incorrectly predicted a cell for a pixel that is background, then there would be a high value for that pixel in the weight map. Alternatively, if the model correctly predicted background for a pixel that was background, then there would be a low value for that pixel in the weight map. This method allows the model to efficiently recognize and discard debris and reduce overfitting on the background. Our code includes a function to automatically generate these weight maps from the ground truth segmentations. Custom weight maps were also implemented for the tracking model, although they were found to be less critical to training a successful model (S6 Fig). The weight map was similarly generated by applying morphological operations to the segmentation of all cells in the current frame and the ground truth. The skeleton of the ground truth cell was set to the highest weight while other cell’s pixels were set to decreasing weights depending on the distance from the tracked cell. Background pixels were all set to a minimal weight of 1. The function generating these maps is also provided in our software.

Two new data augmentations were used while training the segmentation model. We used the blurring function GaussianBlur from the OpenCV package which convolves a 5×5 Gaussian kernel over the image and takes the average of all the pixels under the kernel area. For the noise function we used the random.normal function from the numpy package which outputs random samples from a Gaussian distribution into an array the same size as the image. This is added to the original image and rescaled back to a range between 0 and 1. Both the blurring and noise Gaussian functions are applied to all input images with user-specified standard deviation. We set 1 and 0.03 to be the default standard deviations for the blurring and noise functions, respectively. Another data augmentation operation was used for tracking. To simulate exaggerated cell movement during tracking, such as when an agarose pad dries out and causes the field of view to shift over time, we added a function that randomly shifts the inputs containing the current frame (microscopy image of the current frame, segmentation mask of all the cells in the current frame, ground truth, and weight map) up to five pixels. These operations help expand the training dataset and allow the model to generalize to realistic conditions.

The segmentation model used to quantify the error rate was trained for 600 epochs with 300 steps per epoch and a batch size of 1. The ADAM optimizer was used with a learning rate of 10^-4^. The tracking model used to quantify the error rate was trained for 500 epochs with 300 steps per epoch and a batch size of 2. The ADAM optimizer was used with a learning rate of 10^-5^.

### Training set generation and testing

For the segmentation training dataset, the initial segmentations were generated using the interactive learning and segmentation toolkit Illastik (32). This accounted for 11% of the final training set. Once the segmentation model was performing well on the test data, which we defined as being more than 95% accurate, we used it to generate more training data. Incorrect DeLTA 2.0 outputs, like segmentations that connect two distinct cells together, were manually corrected. Processed DeLTA outputs accounted for 36% of the training set. Additionally, we incorporated published segmentation data from van Vliet *et al*. (33) where cells were segmented and tracked to measure the spatial dynamics of gene expression in bacterial microcolonies. We obtained the cell segmentation and tracking data from the ETH archive: https://doi.org/10.5905/ethz-1007-77. On this dataset, we performed operations to improve the data quality including smoothing filters, dilation, erosion, and skeletonize functions. These data accounted for the remaining 53% of the training set. The final training set had 307 training examples from sixty movies, with a maximum of 10 frames per movie to increase samples diversity. Each training example consisted of a phase contrast image as the input, the corresponding segmented ground truth, and a pre-generated weight map used in the loss function. The test movie used to evaluate the segmentation model was colony 150310-05 from the trpL data zip file from van Vliet *et al*. (33) on the ETH archive.

For the tracking training dataset, we used a modified version of the Matlab script used in Lugagne *et al*. to generate the initial training examples. Instead of showing the whole frame in the graphical user interface, the modified script showed a zoomed-in 75×75 pixel box around the cell of interest. In addition, the modified script had one output consisting of the mother and daughter cell whereas the original script had three outputs for the mother cell, daughter cell, and background. The Matlab script was used to produce 15% of the training set. Each training example consisted of four inputs, one output, and one weight map. The inputs were the phase contrast of the previous frame, segmentation of the cell of interest in the previous frame, phase contrast of the current frame, and segmentation of all the cells in the current frame. The output is a segmentation mask for the cell(s) that the cell of interest in the previous frame tracked to. The weight map is used in the loss function. Once the tracking model was performing on test data with more than 99% accuracy, we used it to generate more training data. Movies with images taken 5 minutes apart were processed using DeLTA 2.0 and then new training examples were generated by tracking cells across longer time intervals. Instead of tracking from the frame immediately before the current timepoint, cells were tracked from a frame from two or three timepoints before. This allowed us to generalize to longer acquisition intervals and to situations where the cells grew faster or travelled further between frames. These processed DeLTA outputs accounted for 20% of the training set. In addition, published tracking data from van Vliet *et al*. was incorporated to increase the training set size, accounting for 65% of all the training examples. The final tracking training set had 23,655 examples. The test movies used to evaluate the tracking model were colony 140408-01 from the cib data zip file; colonies 151029_E1-1, 151029_E1-5, and 151101 _E3-12 from the rpsM data zip file; and colonies 150309-04 and 150310-05 from the trpL data zip file from van Vliet *et al*. (33).

### Time-lapse microscopy experiments

Overnight cultures of *E. coli* MG1655 were diluted 1:100 and allowed to grow for 1-2 hours in LB medium. For the co-culture experiment, we included 30 μg/mL of kanamycin for plasmid maintenance. We created the co-culture with a 1:5 dilution by mixing 0.5 μL of the resistant strain and 0.5 μL of the susceptible strain with 4 μL of LB medium. For the pole age experiment, the culture was diluted 1:100 in MGC media (M9 salts supplemented with 2 mM MgSO4, 0.2% glycerol, 0.01% casamino acids, 0.15 μg/ml biotin, and 1.5 μM thiamine). A 1:100 dilution was used to decrease cell density. For both experiments, 1-1.5 μL of the diluted samples were added to prewarmed 1.5% low melting temperature agarose pads made with MGC media. Samples were prepared and imaged as described in Young, *et al*. (17). A Nikon Ti-E microscope was used with a 100x oil objective for all microscopy experiments.

In the co-culture experiment, *E. coli* MG1655 were transformed with a single plasmid, either with tetracycline resistance or without. Both plasmids originated from the BioBrick plasmid library (pBbA7k) (34). The plasmid for the resistant cells harbored both a tetracycline resistant gene and red fluorescent protein gene (*rfp*) (pBbA7k-RFP-tetA), while the sensitive cells contain only green fluorescent protein (*gfp*) (pBbA7k-sfGFP). The pads contained 30 μg/mL kanamycin for plasmid maintenance and 0.5 μg/mL tetracycline. Phase contrast, GFP, and RFP fluorescence images were taken every 5 minutes.

In the pole age experiment, *E. coli* MG1655 was used, so no antibiotic was present in the culture. Phase contrast images were taken every 5 minutes.

We calculated the growth rate as:

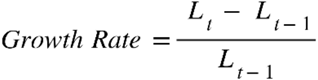

Where *L_t_* is the cell length at time *t*. Growth rates were measured for one generation. For example, to measure the growth rate of OO, measurements start when O divides into OO and end when OO divides into OON and OOO. However, no information about the growth rate of O is used to calculate the growth rate of OO. The growth rate is the average across the timepoints within this generation. To reduce noise, growth rates were only recorded in the analysis if cells were present for at least three frames.

## Acknowledgements

We thank Michael Sheets for testing initial versions of DeLTA 2.0, and members of the Dunlop Lab for helpful discussions. This work was supported by the National Science Foundation grants 2032357 and 1804096 and National Institutes of Health grant AI153853.

## Supporting Information Captions

**S1 Fig. Zoomed out view of Figure 2D.** GFP versus RFP fluorescence for single cells plotted against growth rate. Fluorescence values are the averages over all the frames for that cell. All data points, including outliers, are included in this plot.

**S2 Fig. Limitations of segmentation.** Two sequential phase contrast images of *E. coli* microcolonies with their respective segmentations. The red arrow points to an error where the model has incorrectly combined two cells into one. This type of error is very hard to correct out of context.

**S3 Fig. Reducing the size of the model to increase speed.** Schematics showing different network architectures. **(A)** Original U-Net architecture that we use throughout the paper. **(B)** U-Net architecture without the bottom layer. **(C)** U-Net architecture without the bottom two layers. The model has been trained on segmentation as well as tracking for these reduced networks. The network in (C) runs twice as fast as the original network in (A) and sacrifices very little accuracy.

**S4 Fig. Training the model on segmentation with custom weight maps.** Schematic showing two models trained on the same dataset with different weight maps. In this example, both U-Net models were trained for 600 epochs, 300 steps per epoch, with a batch size of 2. Inputs necessary to train the model include the phase contrast image, the associated segmented ground truth, and weight map. **(A)** Model trained with weight maps derived from the original U-Net paper. Green ovals show examples of errors. **(B)** Model trained with custom weight maps which were generated by applying morphological operations to the segmented ground truth. The model trained on the new weight maps performs better, as shown by the outputs on a test image. **(C)** To aid visualization, the ‘Overlay’ shows the custom weight map overlayed on the phase contrast image. The overlay shows that the weights are emphasized at the core of the cells (shown by red lines) and at the borders (shown by yellow lines).

**S5 Fig. Utilizing variable background weight maps for training the model on debris.** A simplified schematic showing how the loss is calculated for a single input using weight maps. **(A)** Schematic showing the traditional use of weight maps. **(B-C)** Schematics showing the use of variable weight maps. The prediction is used to update the weight map. **(B)** The background weight map values are replaced by the background values in the prediction. This method forces the model to quickly learn debris as the weight map values for the background increase significantly when the model predicts debris as cells. **(C)** Conversely, when the model correctly classifies the debris as background, the weight map values for the background remain similar to the original values.

**S6 Fig. Inputs and outputs required to train the model for tracking.** Left to right, the phase contrast image of the previous frame, segmentation of the cell to be tracked in the previous frame, phase contrast image of the current frame, segmentation of all cells in the current frame, ground truth of the cell being tracked, and weight map. There are four inputs, one output, and one weight map used in the loss function.

**S1 Movie. Time-lapse movie of a bacterial microcolony analyzed with DeLTA 2.0.** Phase contrast images containing *E. coli* cells outlined with different colors representing unique cell numbers. Cells can be tracked by following their respective colors throughout the movie. White arrows indicate cell division events. White and colored dots refer to the new and old poles, respectively. This time-lapse movie was part of the test set used to calculate the error rate for tracking and segmentation. Timestamp shows time in HH:MM.

