## Supplementary figures and images for "DeLTA 2.0: A deep learning pipeline for quantifying single-cell spatial and temporal dynamics"

### Figure S1

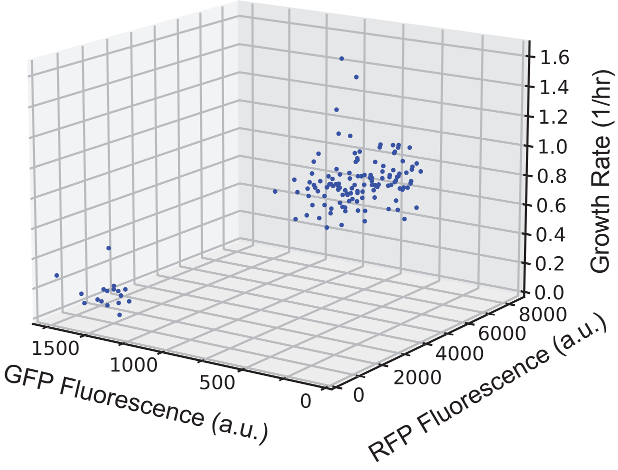

### Figure S2

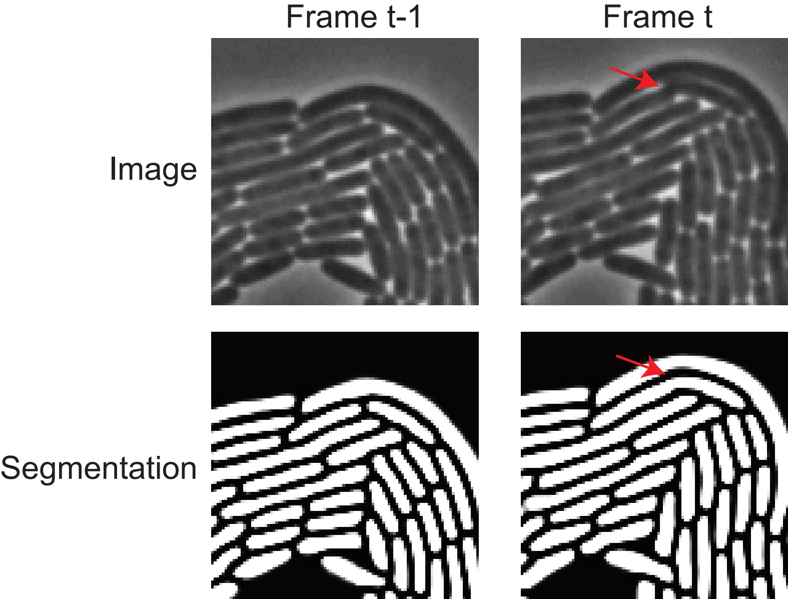

### Figure S3

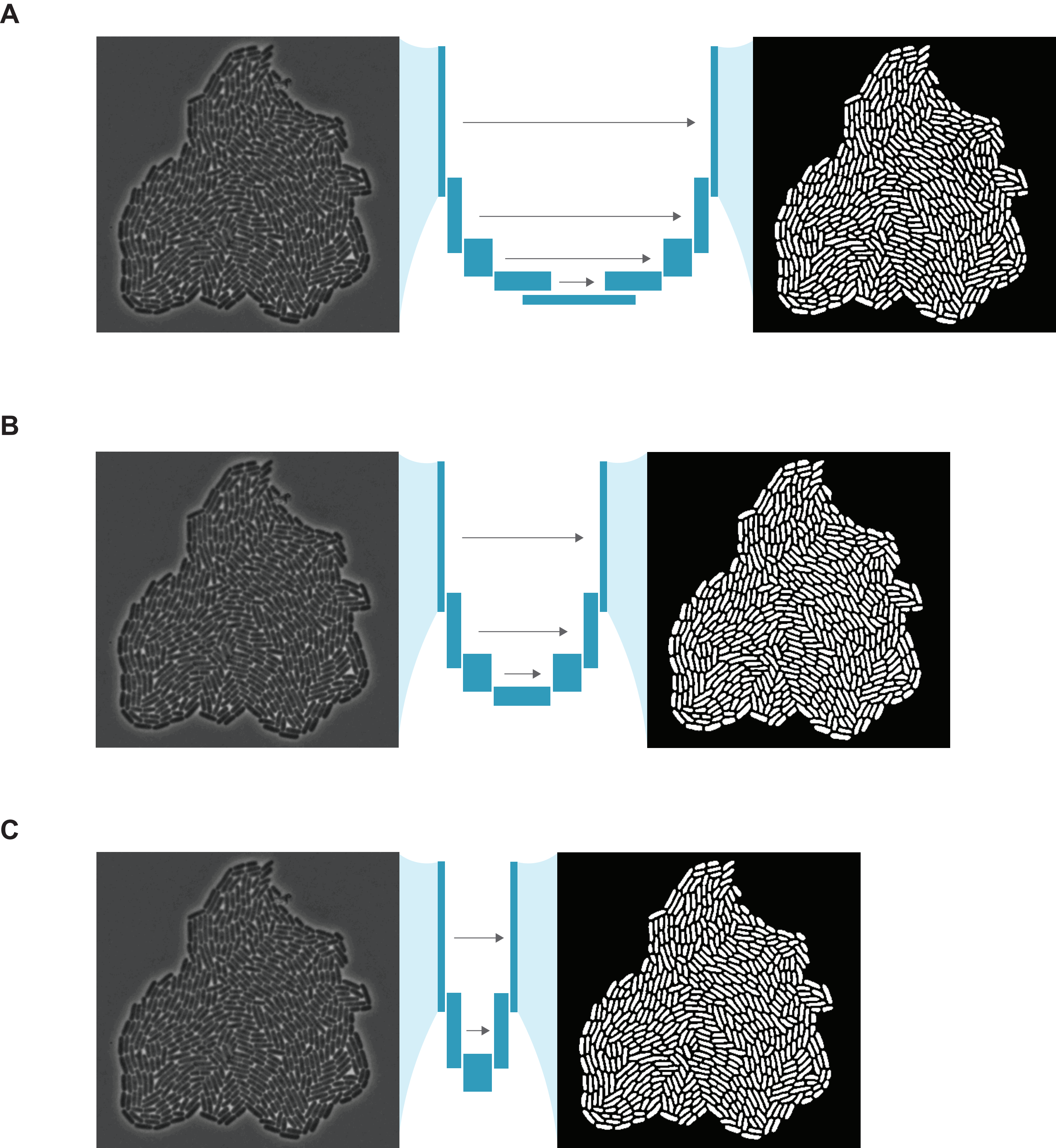

### Figure S4

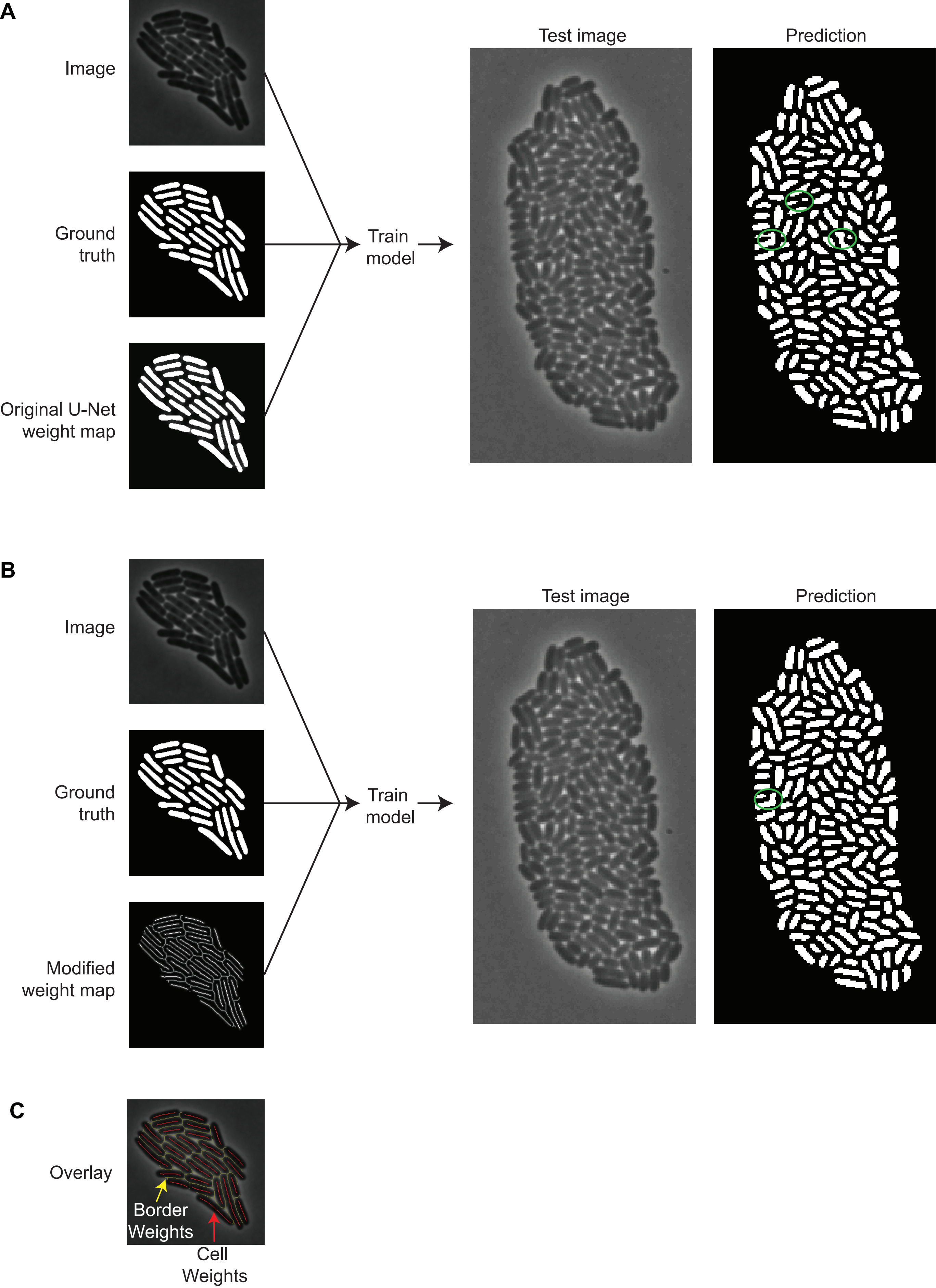

### Figure S5

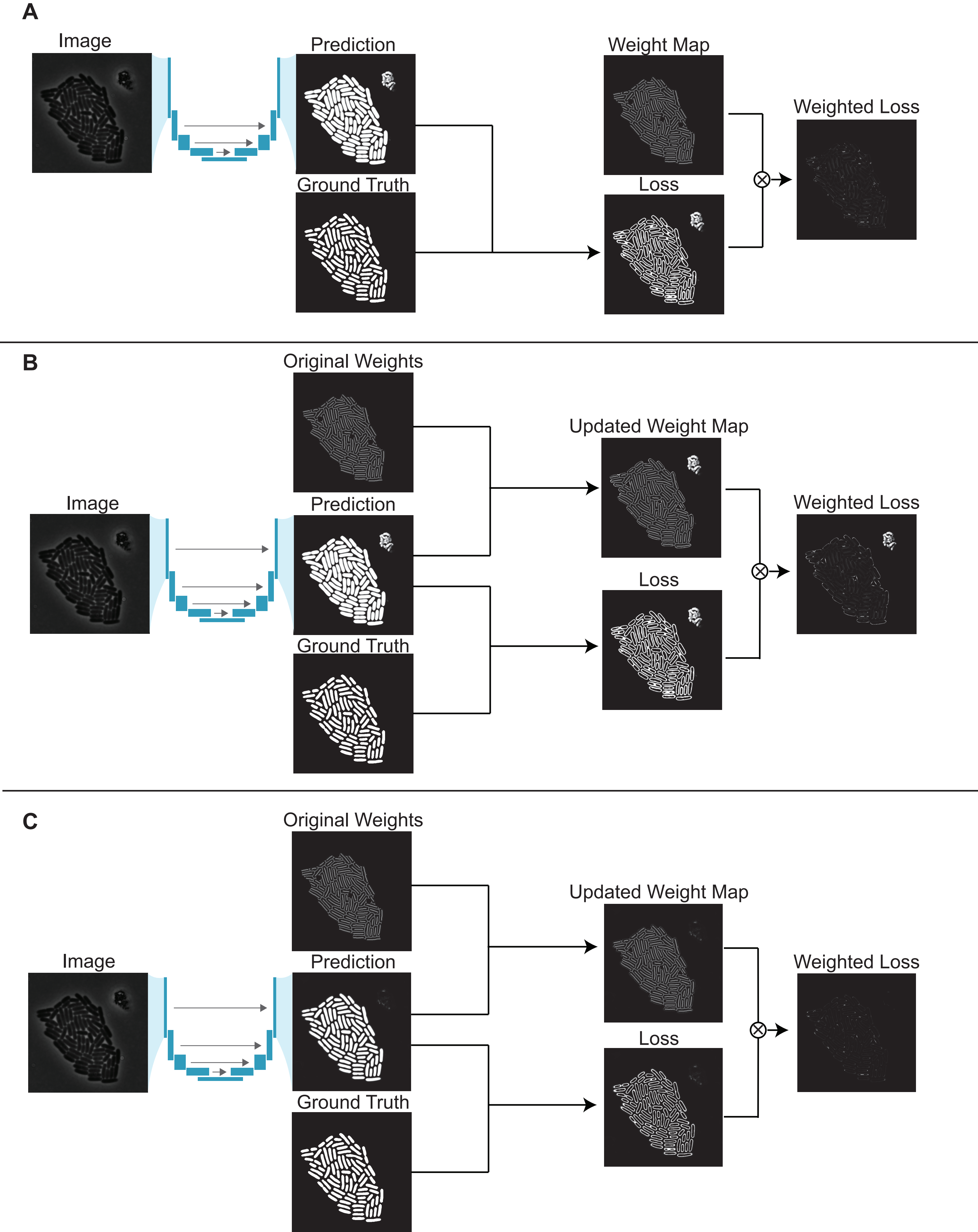

### Figure S6

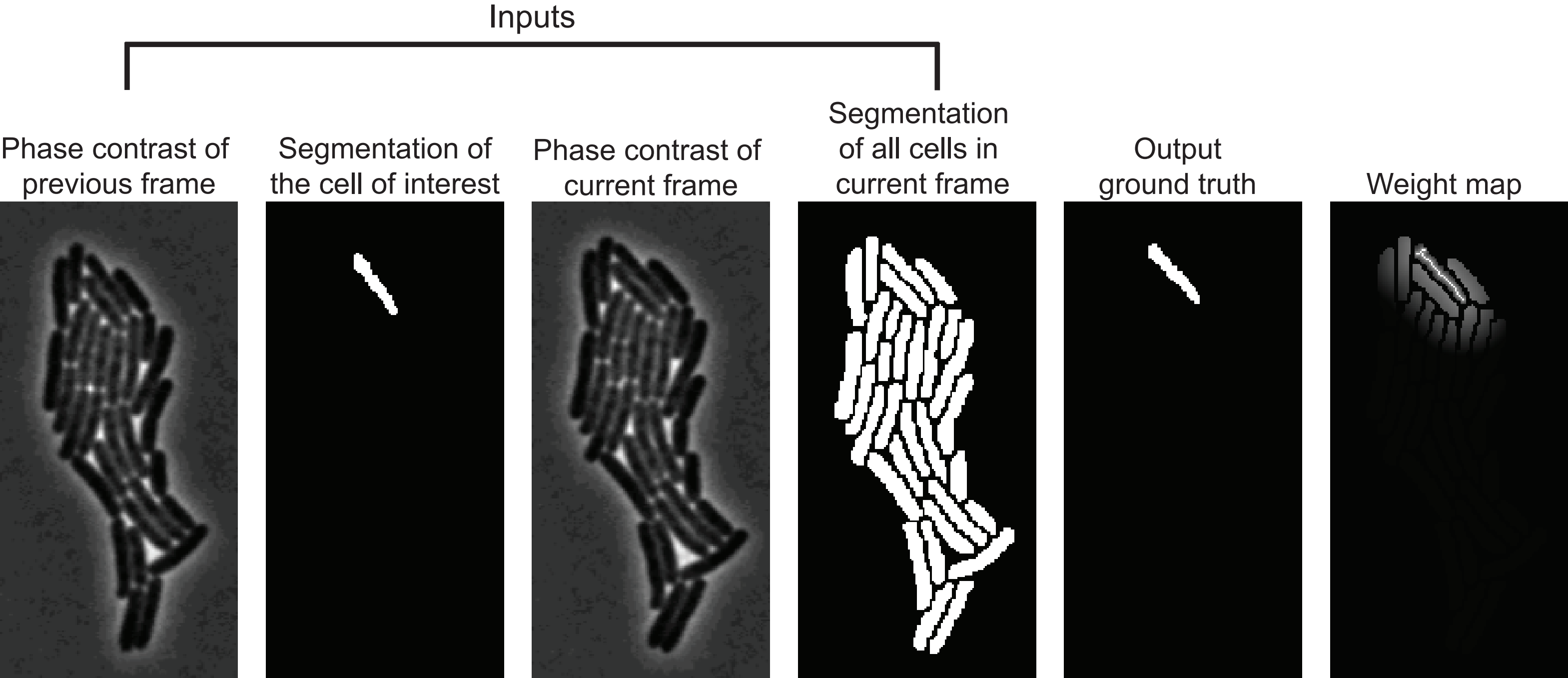
